# Genetic homogeneity in face of morphological heterogeneity in the harbor porpoises from the Black Sea and adjacent waters

**DOI:** 10.1101/634329

**Authors:** Yacine Ben Chehida, Julie Thumloup, Karina Vishnyakova, Pavel Gol’din, Michael C. Fontaine

## Abstract

Isolated from North Atlantic populations, the Black Sea harbor porpoise (*Phocoena phocoena relicta*) is listed as Endangered due to the massive population decline triggered by historical hunting, and subsequently through fisheries bycatch, and other human activities. Of paramount importance for its conservation, is the characterization of the population structure. While morphological heterogeneity suggested population subdivision, previous genetic studies have failed to find any differences. Here, we investigated the population genetic structure of 144 harbor porpoises sampled opportunistically from across the entire subspecies range including the Aegean, Marmara, Black, and Azov Seas. Genetic variation of across one-fourth of the mitochondrial genome, in combination with the analysis of ten microsatellite loci revealed a nearly complete genetic homogeneity. While simulations show that this inability to reject panmixia does not stem from a lack of power (power to detect *F*_*ST*_ of 0.008). A genetic time-lag effect limiting our ability to detect population subdivision is also unlikely when effective population size is low, as is the case here. For now, genetic panmixia among porpoises of the Black Sea and adjacent waters cannot be rejected. Population subdivision may well exist, but conclusive evidence would require an improved sampling providing suitable contrasts (e.g., age, sex, season). Also, a genome scale study providing access to neutral and selected genetic variation may reveal cryptic differentiation indicative of ecologically subdivisions. As a precautionary approach, definition of management units should be based on evidence of population heterogeneity obtained from multidisciplinary approaches rather than just genetics.

## Introduction

Delineating populations and their connectivity is of primary importance for the management of endangered and exploited species (Begg and Waldman, 1999). In marine species, it facilitates identification of stocks, assessing exploitation status, and preserving the population genetic diversity underlying ecological resilience and adaptability (Begg and Waldman, 1999; Palumbi, 2003). Once genetically distinct groups are identified, estimates of their effective size and migration rates is needed to assess their viability and resilience (Frankham, 2010). These population parameters are particularly difficult to estimates for highly mobile species (e.g., marine mammals, turtles, and fishes), yet they are crucially needed to understand the impact of anthropogenic pressures (Payne *et al.*, 2016) and the key roles many of these species play within food webs (Bowen, 1997). The population genetic approach provides a powerful framework for estimating indirectly those parameters (Gagnaire *et al.*, 2015).

Life-history traits of many marine species, such as large population sizes and high dispersal potential, can lead to weak or no genetic differentiation (Waples, 1998; Gagnaire *et al.*, 2015). Indeed, the accumulation of genetic differentiation among populations depends on the effective population size (*Ne*) and the effective number of migrants (*m*) exchanged per generation (*Ne x m*), whereas the level of demographic interdependency depends only on the rate of migrants (*m*) exchanged (Lowe and Allendorf, 2010). In other words, the genetic and demographic connectivity exhibit, in some conditions, a phase difference that prevents the former from being a good proxy of the latter. Such lag is proportional to *Ne*. Common situations involving homogeneous distribution of genetic polymorphism can thus derive from a wide range of distinct demographic scenarios, depending on the relative weight of *Ne* and *m*. These scenarios range from a rate of migratory exchange high enough to lead to both genetic and demographic homogeneity among (sub-)populations, even with limited effective sizes, to nearly negligible migratory exchanges among populations exhibiting large effective sizes. Gagnaire *et al.* (2015) and Bailleul *et al.* (2018) described these effects and showed that the incomplete lineage sorting of populations can be considered as the homologous version at an intraspecific level of the “*grey zone*” of speciation described by De Queiroz (2007). This “*grey zone*” represents the lag during which, lineage sorting being incomplete, species delimitation is not possible based solely on the genetic information (Gagnaire *et al.* 2015; Bailleul *et al.*, 2018). This concept of “*grey zone"* of population differentiation was coined by Bailleul *et al.* (2018) as the number of generations after a population split for genetic drift to change the allele frequencies in each diverging population and reach an equilibrium between migration and genetic drift (Epps and Keyghobadi, 2015). During that period, a time-lag between genetic and demographic structure occurs, and no decision can be made from genetic data to assess whether two groups are demographically independents based solely on genetic data. The length of that period increases with *Ne*. It is therefore critical to assess whether the lack of genetic structure observed in a particular case results from such a time-lag effect, from a lack of genetic power, or from an actual demographic and genetic homogeneity. Marine species with large *Ne*, high fecundity, and high dispersal abilities, such as fishes or invertebrates, are the primary species where such lag between genetic and demographic processes is expected (see for example (Waples, 1998; Palumbi, 2003; Gagnaire *et al.* 2015)). In contrast, theory predicts that species with smaller *Ne*, lower fecundity, but high dispersal abilities, such as marine mammals, should have a narrower population *grey zone*. Observing a genetic panmixia in those species is thus more likely to reflect an actual absence of genetic and demographic population structure, rather than a demographic independence not yet captured by genetic data.

The subspecies of harbor porpoises inhabiting the Black Sea (*Phocoena phocoena relicta*) is a good example to illustrate this point. The harbor porpoise is one of the three extant cetacean species crowning the Black Sea marine trophic food-web. *P. p. relicta* became isolated ca. 7,000 years ago from the rest of the species range in the North Atlantic during the postglacial warming of the Mediterranean Sea, which became unsuitable for temperate species like porpoises (Fontaine *et al.* 2010; 2012; 2014 and reviewed in Fontaine, 2016). Black Sea porpoises are recognized as a distinct subspecies based on morphological and genetic differences as compared to the North Atlantic porpoises (*P. p. phocoena*) (Viaud-Martinez *et al.*, 2007; Fontaine *et al.*, 2007; 2014; Galatius and Gol’din, 2011; and reviewed in Fontaine, 2016). In the Black Sea and adjacent waters (Fig. 1), porpoises are observed in the northern Aegean Sea, Marmara Sea, Black Sea, Kerch Strait and Azov Sea (Fontaine, 2016). The Black Sea harbor porpoise is listed as Endangered by the IUCN (Birkun and Frantzis, 2008). These porpoises have been hunt near the extinction level between from 1930’s to 1980’s, causing a ~90% population decline (Birkun, 2002; Fontaine *et al.*, 2012; Vishnyakova, 2017). Subsequent incidental catches in fisheries reached thousands of porpoise casualties annually through the 1980’s and are likely to have increase since then (Birkun and Frantzis, 2008; Vishnyakova and Gol’din, 2015; Vishnyakova, 2017). Major mass mortality event occurred in the Azov Sea in August 1982 as a result of an explosion at a gas-extraction platform killing more than 2,000 porpoises. Other mass mortalities were reported, some attributed to diseases or ice entrapments, and possibly also to habitat degradation including prey’s stock depletion, pollution, eutrophication, noise disturbance, etc. (Birkun and Frantzis, 2008; Daskalov, 2003; Vishnyakova and Gol’din, 2015; Vishnyakova, 2017). Having a clear picture of the genetic structure is thus crucial for devising conservation strategies (Allendorf *et al.*, 2012).

**Fig. 1.**
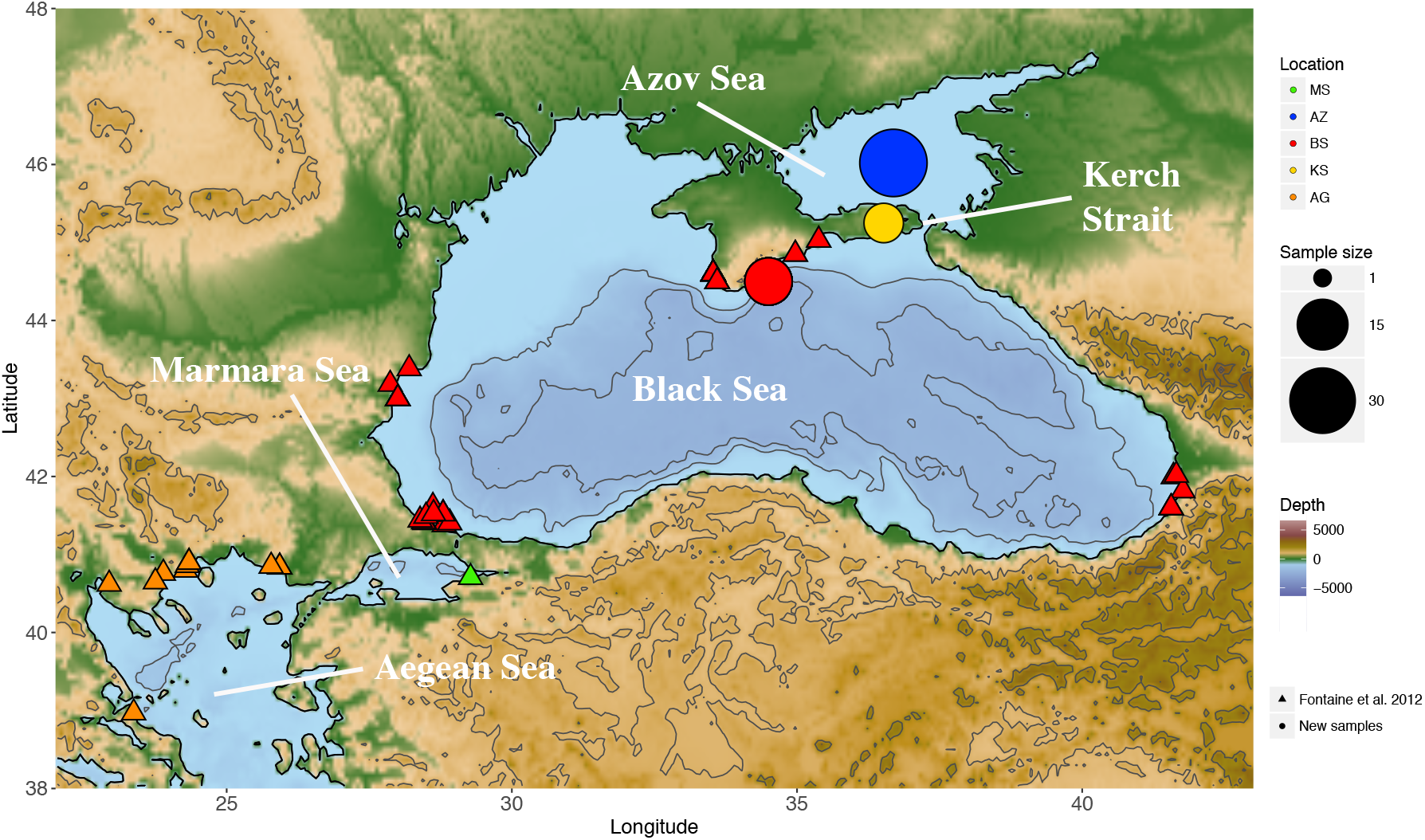
Map showing the sampling locations. The radius of the circles is proportional to the sampling size. AG, Aegean Sea; MS, Marmara Sea; BS, Black Sea; KS, Kerch Strait; AZ, Azov Sea. Rectangles and circles represent individuals sampled respectively from Fontaine *et al.* (2012) and this study.

It is still unclear whether the Black Sea porpoises are composed of a single homogeneous demographic and genetic unit or multiple interconnected ones, but differentiated enough to be considered as distinct populations. Some authors suggested that population subdivision might exist (Rosel *et al.*, 2003; Gol’din, 2004). For example, Rosel *et al.* (2003) hypothesized that porpoises from the Aegean and Black Seas could belong to distinct subpopulations. Genetic analyses of the mitochondrial control-region lent some support for this hypothesis with subtle but statistically significant differences (Viaud-Martinez *et al.*, 2007) or between those from the Marmara and Black Seas (Tonay *et al.*, 2017). However, these observations were based on a small sample size and a single locus. In contrast, other studies combining highly polymorphic nuclear microsatellites and mitochondrial loci failed to detect such genetic structure and suggested that these porpoises were panmictic (Fontaine *et al.*, 2012). Similarly, morphological differences between porpoises from the Black and Azov Seas suggested that they may be differentiated subpopulations (Gol’din, 2004; Gol’din and Vishnyakova, 2015, 2016). For instance, compared to animals from the Black Sea, porpoises from the Azov Sea display slightly larger body size (Gol’din, 2004) and distinct skull sizes and shapes (Gol’din and Vishnyakova, 2015, 2016). The authors suggested that these differences may reflect distinct feeding ecology, ontogeny, and thus possibly demographically distinct units. However, so far, no genetic analysis has tested these hypotheses, and none included the porpoises from the Azov Sea.

This study aims at providing a global picture of the genetic structure of the harbor porpoises in the Black Sea and adjacent waters. Following on Fontaine *et al.* (2012), we expanded the previous genetic data set (utilizing autosomal microsatellite and mitochondrial loci) with 55 new samples from the Black Sea, the Sea of Azov, and the Kerch Strait (Fig. 1). Specifically, we tested whether the evidence of population subdivision previously reported based on phenotypic or mitochondrial variation can actually be recovered with multiple independent genetic loci that have distinct but complementary inheritance modes (maternally inherited mitochondrial locus *versus* biparentally inherited autosomal microsatellite loci). We used a simulation framework to test whether an absence of genetic structure results from limited power or to a lag between genetic and demographic signal.

## Materials and Methods

### Sampling and data collection

The samples used in this study originated from 5 geographic locations: the Aegean Sea, Marmara Sea, Black Sea, Kerch Strait, and Azov Sea (Fig. 1 and Table 1). Genotypes at 10 microsatellite loci for 89 porpoises from the Aegean, Marmara, and Black Seas were taken from Fontaine *et al.* (2012). We added 55 newly genotyped individuals from the Azov Sea, Kerch Strait, and Black Sea (Fig. 1 and Table 1). Tissue samples were collected from dead animals stranded along the coasts of the Crimea (Ukraine) on the Black Sea, Kerch Strait, and the Azov Sea and kept in DMSO until analyses. Total genomic DNA was extracted from tissues using a PureGene and DNeasy Tissue kit (Qiagen), following the manufacturer’s recommendations. The microsatellite genotyping procedure followed the protocol described in Fontaine *et al.* (2006; 2007).

**Table 1.**
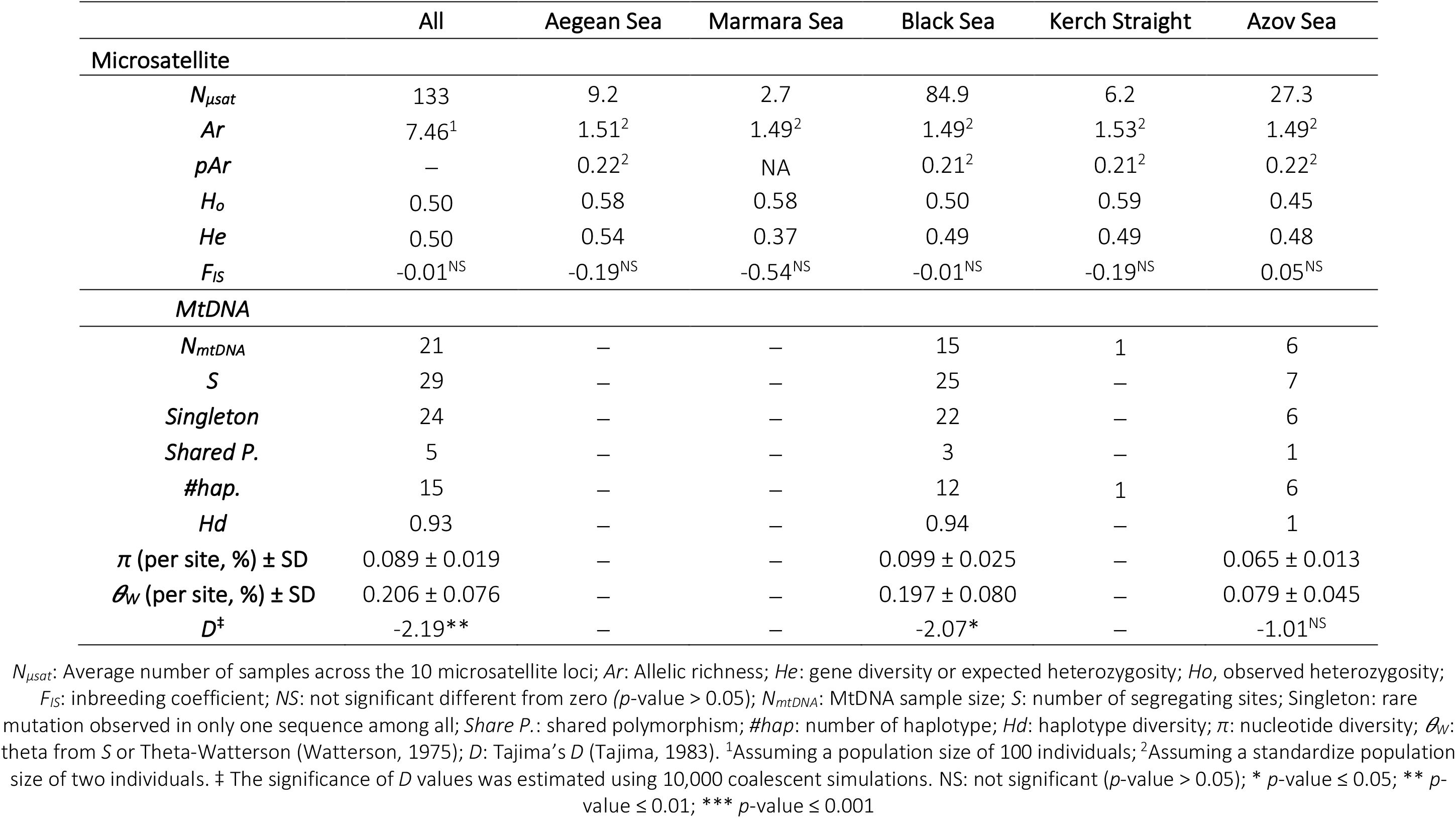
Genetic diversity at the 10 microsatellites loci and mitochondrial locus.

|  | All | Aegean Sea | Marmara Sea | Black Sea | Kerch Strait | Azov Sea |
| --- | --- | --- | --- | --- | --- | --- |
| **Microsatellite** |  |  |  |  |  |  |
| $N_{\mu sat}$ | 133 | 9.2 | 2.7 | 84.9 | 6.2 | 27.3 |
| $Ar$ | 7.46<sup>1</sup> | 1.51<sup>2</sup> | 1.49<sup>2</sup> | 1.49<sup>2</sup> | 1.53<sup>2</sup> | 1.49<sup>2</sup> |
| $pAr$ | — | 0.22<sup>2</sup> | NA | 0.21<sup>2</sup> | 0.21<sup>2</sup> | 0.22<sup>2</sup> |
| $H_o$ | 0.50 | 0.58 | 0.58 | 0.50 | 0.59 | 0.45 |
| $He$ | 0.50 | 0.54 | 0.37 | 0.49 | 0.49 | 0.48 |
| $F_{IS}$ | -0.01<sup>NS</sup> | -0.19<sup>NS</sup> | -0.54<sup>NS</sup> | -0.01<sup>NS</sup> | -0.19<sup>NS</sup> | 0.05<sup>NS</sup> |
| **MtDNA** |  |  |  |  |  |  |
| $N_{mtDNA}$ | 21 | — | — | 15 | 1 | 6 |
| $S$ | 29 | — | — | 25 | — | 7 |
| *Singleton* | 24 | — | — | 22 | — | 6 |
| *Shared P.* | 5 | — | — | 3 | — | 1 |
| *#hap.* | 15 | — | — | 12 | 1 | 6 |
| $Hd$ | 0.93 | — | — | 0.94 | — | 1 |
| $\pi$ (per site, %) ± SD | 0.089 ± 0.019 | — | — | 0.099 ± 0.025 | — | 0.065 ± 0.013 |
| $\theta_w$ (per site, %) ± SD | 0.206 ± 0.076 | — | — | 0.197 ± 0.080 | — | 0.079 ± 0.045 |
| $D^\dagger$ | -2.19** | — | — | -2.07* | — | -1.01<sup>NS</sup> |

$N_{\mu sat}$ : Average number of samples across the 10 microsatellite loci; $Ar$ : Allelic richness; $He$ : gene diversity or expected heterozygosity; $H_o$ , observed heterozygosity; $F_{IS}$ : inbreeding coefficient; NS: not significant different from zero ( $p$ -value > 0.05); $N_{mtDNA}$ : MtDNA sample size; $S$ : number of segregating sites; Singleton: rare mutation observed in only one sequence among all; *Share P.*: shared polymorphism; *#hap.*: number of haplotype; $Hd$ : haplotype diversity; $\pi$ : nucleotide diversity; $\theta_w$ : theta from $S$ or Theta-Watterson (Watterson, 1975); $D$ : Tajima's $D$ (Tajima, 1983). <sup>1</sup>Assuming a population size of 100 individuals; <sup>2</sup>Assuming a standardize population size of two individuals. † The significance of $D$ values was estimated using 10,000 coalescent simulations. NS: not significant ( $p$ -value > 0.05); \* $p$ -value ≤ 0.05; \*\* $p$ -value ≤ 0.01; \*\*\* $p$ -value ≤ 0.001

In addition, we sequenced a 3,904 base-pair fragment of the mtDNA genome encompassing five coding regions (CytB, ATP6, ATP8, ND5 and COXI) for 10 individuals (Azov Sea: N=6, Black Sea: N=3 and Kerch Strait: N=1) and combined it with the 12 sequences previously obtained for porpoises from the Black Sea in Fontaine *et al.* (2014), following the same protocol. We used *Geneious* v.10.0.9 (Kearse *et al.*, 2012) to visually inspect raw sequences, assemble contigs, and perform multiple sequences alignments using MUSCLE (Edgar, 2004).

### Genetic diversity among geographic locations

The genetic diversity at the microsatellite loci was quantified over the entire sampling (global) and per geographic location (local) using the allelic richness (*Ar*), expected heterozygosity (*He*) and observed heterozygosity (*Ho*). Global *Ar* was calculated using *Fstat* v.2.9.3.2 (Goudet, 1995). Global and local *Ho* and *He* were calculated using *GenAlEx* v.6.5 (Peakall and Smouse, 2012). Local *Ar* and private *Ar* (*PAr*) were estimated using *ADZE* (Szpiech *et al.*, 2008) assuming a standardized sample size of 2 individuals. We tested for significant differences in *Ar*, *Par*, *Ho* and *He* among locations using a Wilcoxon signed-ranked tests. Adjustment for multiple comparisons was performed using a Bonferroni correction (error rate α = 0.05). Overall departure from Hardy Weinberg Expectation (HWE) was tested using an exact test (Guo and Thompson, 1992) implemented in *Genepop* v.4.7.0 (Rousset, 2008) and quantified this departure using the *F*_*IS*_ estimator of Weir and Cockerham (1984) in *GenAlEx* v.6.5.

The variation among mitochondrial sequences was assessed using various statistics including the number of segregating sites (*S*), number of singletons, number of shared polymorphisms (*Shared P.*), number of haplotypes (*#hap.*), haplotype diversity (*H*_*d*_), two estimators of population genetic diversity *𝛝*_*π*_ (Tajima, 1983) based on the average number of pairwise differences (*K*), and *𝛝*_*w*_ (Watterson, 1975) based on the number of segregating sites. All these statistics were computed using *DnaSP* v.5.10.01 (Librado and Rozas, 2009).

Phylogenetic relationships among mtDNA haplotypes were estimated using the maximum-likelihood approach of *PhyML* v3.0 (Guindon *et al.*, 2010) implemented as a plug-in in *Geneious* v.10.0.9 (Kearse *et al.*, 2012). We used *jModelTest2* (Darriba *et al.*, 2012) to select the model of nucleotide substitution best fitting with our sequence alignment. The tree was rooted with two mitochondrial sequences of Dall’s porpoise from Fontaine *et al*. (2014). We draw the phylogenetic trees using *FigTree* v.1.4.3 (Rambaut and Drummond, 2012). Node support was estimated using 1 × 10^4^ bootstrap replicates. As a complementary visualization of phylogenetic relationships among haplotypes, we also reconstructed a Median-Joining haplotype network (Bandelt *et al.*, 1999) using PopART (http://popart.otago.ac.nz).

### Relatedness

Considering closely related individuals to delineate population genetic structure can generate spurious signal of population structure and violates the assumptions of population genetic approaches, such as the model-based Bayesian clustering (Anderson and Dunham, 2008; RodriguezRamilio and Wang, 2012). Therefore, we analyzed patterns of relatedness among individuals using the *R* package *related* v.1.0 (Pew *et al.*, 2015) in the R statistical environment v.3.5.3 (R Core Team, 2019). Specifically, we estimated the relatedness coefficient (*r*) among individuals and tested whether it was greater within each location than expected by chance. As the performance depends on the characteristics of the data set (Csilléry *et al.*, 2006) and on the estimators, we compared seven estimators implemented in the package *related* following the user-guide recommendation. We identified the Wang estimator (Wang, 2002) as the estimator providing the best performance for our dataset (results not shown). We assessed whether individuals within each location are more closely related to each other than expected by chance. To do so, we compared the observed *r* value in each location against the null distribution of individuals pairwise average *r* generated by randomly shuffling individuals among populations for 1000 permutations while keeping the population size constant. An empirical *p*-value was obtained by comparing the observed average *r* for each population with the null distribution by counting the number of times the observed *r* was greater than those obtained from permuted data. We applied a Bonferroni correction to adjusted for multiple comparisons with a significance threshold of 0.01.

### Population genetic structure

We assessed the genetic structure among porpoises with the Bayesian clustering approach of *STRUCTURE* v.2.3.4 (Pritchard *et al.*, 2000; Hubisz *et al.*, 2009) using an admixture “*locprior*” model and correlated allele frequencies among clusters (Hubisz *et al.*, 2009). This parametrization is suitable for detecting weak genetic structure when it exists, yet without forcing it (Hubisz *et al.*, 2009). The sampling location of each individual was used as *prior* information in the *locpior* model. We conducted a series of independent runs with different number of clusters (*K*) ranging from 1 to 7. Each run used 1 × 10^6^ iterations after a burn-in of 1×10^5^ iterations with 10 replicates per *K* value. We assessed convergence of the Monte Carlo Markov Chains (MCMC) using *CLUMPAK* (Kopelman *et al.*, 2015). We determined the best *K* value using (1) the log likelihood of the data for each K value, (2) the rate of change of K with increasing *K* (Evanno *et al.*, 2005), and (3) the visual inspection of newly created cluster as K increased. For the step (1) and (2) we used *STRUCTURE HARVESTER* v.0.6.94 (Earl and vonHoldt, 2011).

We also investigated the genetic structure using a Principal Component Analysis (PCA) on the allele frequencies (Jombart *et al.*, 2009). This analysis does not rely on any model assumptions and provides a complementary visualization of the genetic structure. This analysis was conducted in R (R Core Team, 2019) using the package *adegenet* (Jombart, 2008; Jombart and Ahmed, 2011) on centered data, with missing data replaced by the mean. We also conducted a Discriminant Analysis of Principal Components (DAPC) (Jombart *et al.*, 2010). The analysis uses the principal components of the PCA to maximizes differences among *predefined* groups using a discriminant analysis. We used the sampling locations as putative grouping. The number of PCs retained and the reliability of the DAPC were assessed using the *a*-score approach following the user guide recommendation. As a result, a total of 21 PCs and 4 discriminant functions were retained to describe the relationship between the clusters, which captured 91% of the total genetic variation.

### Genetic differentiation among populations

For microsatellites data, we estimated the overall and pair-wise departure from HWE due to population subdivision using the *F*_*ST*_’s Weir and Cockerham estimator (Weir and Cockerham, 1984). The *F*_*ST*_ 95% confidence interval (95% CI) was estimated using 5000 bootstrap resampling with the *R* package *DiveRsity* v1.9.90 (Keenan *et al.*, 2013). The significance was tested using an exact *G*-test (Goudet *et al.*, 1996) implemented in *Genepop* v.4.7.0 (Rousset, 2008) with default options. We used a Bonferroni correction to adjust the *p*-value to 0.05 of the pair-wise comparisons to account for multiple comparisons. For mtDNA data, we quantified the genetic differentiation between the Black Sea and Azov Sea using the Hudson’s estimator of *F*_*ST*_ (Hudson *et al.*, 1992) using *DnaSP* v.5.10.01. Significance was tested with 10,000 permutations of Hudson’s nearest neighbour distance *Snn* statistics (Hudson, 2000) in *DnaSP* v.5.10.01.

We assessed the statistical power of our markers to detect genetic differentiation given the observed markers diversity and sample sizes using the program *POWSIM* v4.1 (Ryman and Palm, 2006). *POWSIM* assesses whether the observed data set carry enough statistical power (*i.e. ≥*80%) to detect a Nei’s *F*_*ST*_ (*F*_*ST-Nei*_) value significantly larger than zero using Chi^2^ and Fisher tests (Ryman and Palm, 2006). Parameters of the Markov chains, including the burn-ins, batches and iterations per run were set respectively to 10000, 200, and 5000. Alleles frequencies were estimated with *GenAlEx* and haplotypes frequencies with *DnaSP*. Sample sizes were adjusted for the mtDNA to reflect the sampling of haploid genes (Larsson *et al.*, 2008). Observed *F*_*ST-Nei*_ for microsatellite and mitochondrial data were calculated using *DiveRsity* v1.9.90 (Keenan *et al.*, 2013), and the R package *mmod* v.1.3.3 (Winter, 2012), respectively. *Ne* was fixed to 1,000 and the number of generations (*t*) was adjust to obtain *F*_*ST-Nei*_ values ranging from 0.001 to 0.15 for microsatellites and from 0.001 to 0.4 for mtDNA.

### Simulations of population connectivity and “*grey zone*” of population differentiation

We assessed whether an absence of significant genetic structure could result from a time lag effect between demographic and genetic processes using the same simulation approach of Bailleul *et al.* (2018) adapted to our system. Simulations are used to assess the number of generations required to overcome the “population *grey zone*” and be able to detect *F*_*ST*_ values significantly greater than 0 for a pair of diverging populations. Specifically, we used *simuPOP* v.1.1.7 (Peng and Amos, 2008) to conduct forward-time simulations of two diverging populations of random mating individuals (recombination rate of 0.01) to generate genetic data sets with similar properties as the observed one (10 loci with 10 allelic states). To mimic the founding event of the Black Sea subspecies 700 generations ago (or *ca.* 7000 years before present) (Fontaine *et al.*, 2010; 2012; 2014), we simulated an initial population with an effective size *Ne*_*ini*_, that split 700 generations ago into two daughter populations, each diverging from each other with a constant effective population size *Ne*_*cur*_. As the time to overcome the population “*grey zone*” depends on *Ne* and *m*, we ran the simulations assuming three values for *Ne*_*ini*_ = 10, 100 or 1,000, testing thus a gradient in the strength of the founding effect, one value of *Ne*_*cur*_ = 1,000, and three values of symmetrical migration rates *m* set in such a way that the effective number of migrants per generation (*Ne*_*cur*_ x *m*) is equal to 0 (no migration), 1, 10 or 100. For each of the 12 parameter combinations, we sampled the *F*_*ST*_ values during the differentiation process every 7 generations (100 data points). For each time point, *F*_*ST*_ value were estimated based on the 1000 individuals in each population. Furthermore, at each time point, 100 sub-*F*_*ST*_ values were estimated based on a realistic subsample of 50 in each diverging population. Significance of the sub-*F*_*ST*_ was assessed by randomly shuffling 1000 times the individuals in the subsamples and computing an *F*_*ST*_. A *p-value* was derived from this null randomized *F*_*ST*_ distribution and estimated as the proportion of randomized *F*_*ST*_ inferior or equal to the simulated sub-*F*_*ST*_. Finally, our ability to detect a *F*_*ST*_ value significantly greater than 0 (in percent) was estimated by counting, out of the 100 replicates, the proportion of sub-*F*_*ST*_ with *p-values* ≤ 0.05.

## Results

### Genetic diversity

Out of the 144 individuals genotyped for the 10 microsatellite loci, the level observed missing data reached a total of 6.93%. Across all geographic areas and loci (Table 1 and S1), we observed an allelic richness (*Ar*) of 7.5 and a genetic diversity (*He)* of 0.50. No departure from Hardy-Weinberg Equilibrium (HWE) was observed (*F*_*IS*_ = −0.01, *p*-value = 0.956), indicating no detectable departure from panmixia. Also, we observed a genetic homogeneity in genetic diversity across the 5 sampled areas, each one displaying no deviation from HWE and no detectable differences in their genetic diversity (Table 1 and S1). For a standardized sample size of 2 individuals, *Ar* and private *Ar* (*pAr*) values ranged from 1.49 to 1.51 and from 0.21 to 0.30 among the 5 sampled areas respectively, without any significant difference among them (Table 1; Wilcoxon signed-ranked (WSR) test with a *p-value* > 0.05). The expected and observed heterozygosity (*Ho* and *He*) were also comparable among geographic areas, ranging between 0.45 and 0.59 (WSR test with a *p-value* > 0.05).

For the 3,904bps mtDNA fragment analyzed, a total of 25 segregating sites defined 15 distinct haplotypes with a haplotypic diversity of 0.93 ± 0.05 and a nucleotide diversity of 0.089% ± 0.019% (Table 1). The phylogenetic relationships among haplotypes revealed a star-like topology on the maximum likelihood phylogenetic tree (Fig. 2a) and haplotype network (Fig. 2b) where rare haplotypes are all closely related to a dominant haplotype, with only one or two distinct mutations. This topology is consistent with significant excess of rare variant over the shared variant as capture by significant negative value of Tajima’s *D* statistics (−2.19; P < 0.01; Table 1). Out of 15 haplotypes, three were unique to the porpoises from the Azov Sea, nine only found in the Black Sea porpoises, and three were shared between the two (Table 1, Fig. 2a and 2b). There was no obvious clustering according to the geography.

**Fig. 2.**
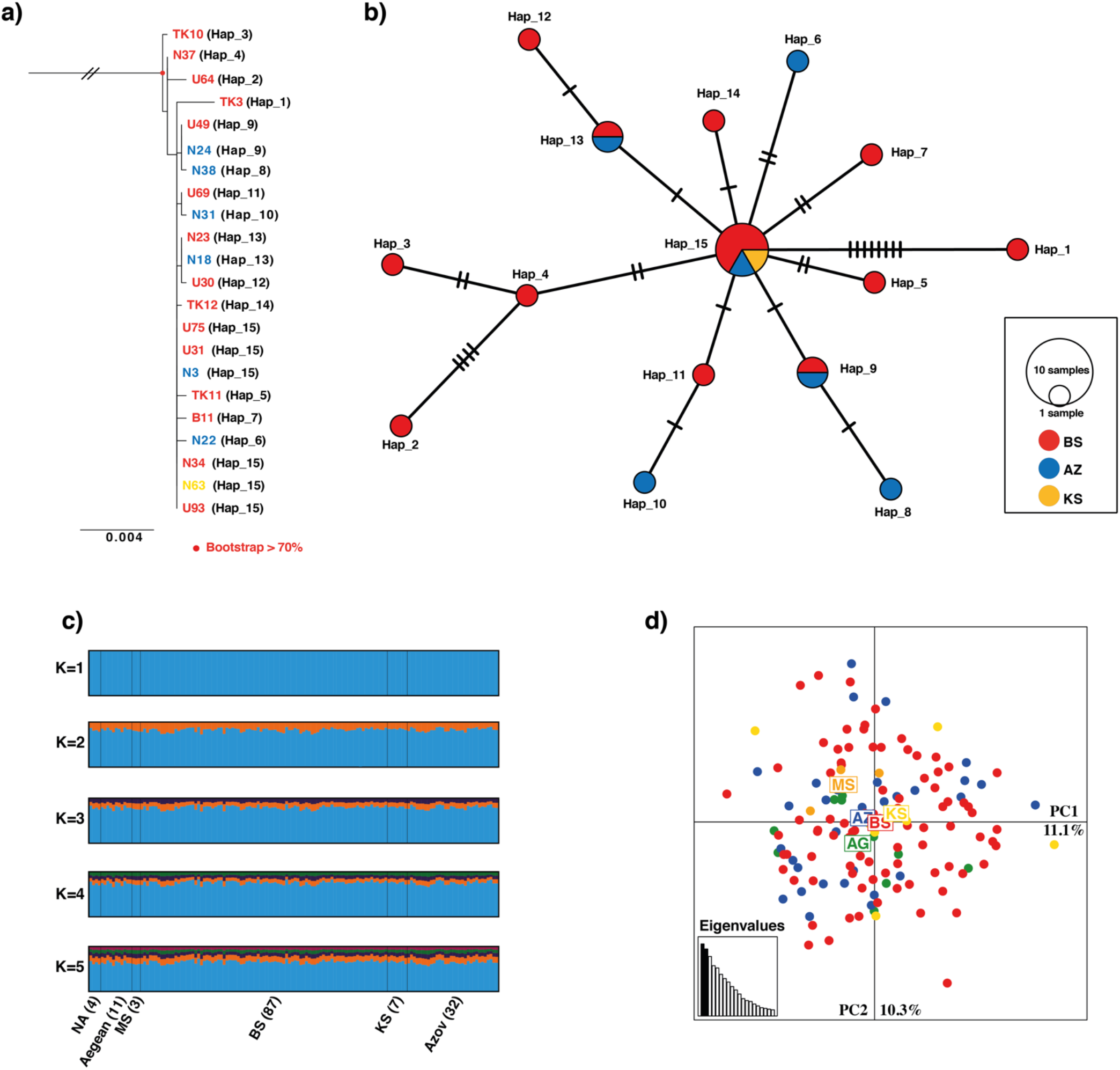
Population structure observed at the mtDNA and microsatellite loci. a) Maximum-likelihood mitochondrial phylogeny rooted with Dall’s porpoise sequences (not shown). The labels’ colors indicate the sampling location (blue: Azov Sea or AZ; red: Black Sea or BS; yellow: Kerch Strait or KS). Nodes’ color in red represent the bootstrap supports >70%. b) Median-joining mitochondrial haplotype network. Each circle represents a haplotype and the size is proportional to observed haplotype frequency. Pie-chart sectors indicate the number of haplotypes observed in each locality. Mutational steps between haplotype are represented on the branch. c) Barplots of the Bayesian clustering analyses of *STRUCTURE* for K from 1 to 5. Each individual is represented by a vertical line divided into K segments showing the admixture proportions to each cluster. Vertical black lines delimit the sampled localities. d) Scatter plot displaying the individual scores along the first two components of the principal component analysis. The proportion of variance explained by each axis and the first eigenvalues (bottom left inset) are provided. AG, Aegean Sea; MS, Marmara Sea; BS, Black Sea; KS, Kerch Strait; AZ, Azov Sea.

### Relatedness

Relatedness estimates (*r*) among porpoises within each sampled locality revealed that only the three individuals from the Marmara Sea displayed an *r* value much greater than expected by chance alone (*p*-value < 0.001). The average *r* for these individuals was 0.55 ± 0.15, which correspond to a parent-offspring or full sibling relationship. For all other populations, *r* values ranged from 0.06 to 0.08 as expected for unrelated individuals (Fig. S1 and Table S2).

### Genetic structure

The clustering analyses of STRUCTURE did not reveal any evidence of population subdivision irrespective of the number clusters *K* tested (Fig. 2c and Fig. S2a). The highest posterior probability for the data (X) of containing *K* clusters Ln(*Pr(X|K*) was observed for *K*=1 and *K*=2 and was much lower for higher *K* values (Fig. S2b). Regardless of the *K* value tested, individual pattern of admixture was identical for all individuals, suggesting that harbor porpoises from the different localities behave as a panmictic population. The analysis provided consistent results over 10 replicated runs performed for each *K* (Fig. S2a).

The principal component analysis (PCA) supported the results of *STRUCTURE* by showing no evidence of population subdivision, as all multilocus genotypes group into a unique cluster (Fig. 2d). The Discriminant Analysis of Principal Component (DAPC; Fig. S3), which focus on optimizing the differences between predefined clusters, here the sampled localities, while minimizing the differences within groups, showed globally similar results as *STRUCTURE* (Fig. 2c). No genetic subdivision could be observed between individuals from the Black Sea and the Azov Sea which are located in the center of the DAPC (Fig. S3). Similarly, there is no clear separation between the individuals from the Kerch Strait and Aegean Sea. Only the second discriminant function slightly discriminates the porpoises from the Marmara Sea from the others (Fig. S3), very likely as a result of their high relatedness (Anderson and Dunham, 2008; RodriguezRamilio and Wang, 2012).

The absence of genetic structure at the microsatellite loci was further supported by the very low global *F*_*ST*_ values (*F*_*ST-WC*_ = 0.009 and Nei’s *F*_*ST-Nei*_ = 0.022), not significantly departing from zero (*p*-value=0.109). Similarly, all pairwise comparisons displayed non-significant differences in allelic frequencies (Table 2). Only the *F*_*ST*_ values between Marmara and Aegean porpoises was slightly higher (*F*_*ST-WC*_ ≥ 0.044 and *F*_*ST-Nei*_ ≥ 0.017), but none departed significantly from zero, and only the *F*_*ST-WC*_ between Aegean Sea and the Marmara Sea did not include 0 in the 95% CI (Table 2). We did not detect any signal of population differentiation at the mtDNA locus (Hudson’s *F*_*ST*_ = 0.007, Nei’s F_ST_=0.013, Snn=0.519, Snn’s *p*-value=0.726) suggesting no genetic subdivision between the porpoises from the Azov and Black Seas.

**Table 2:**
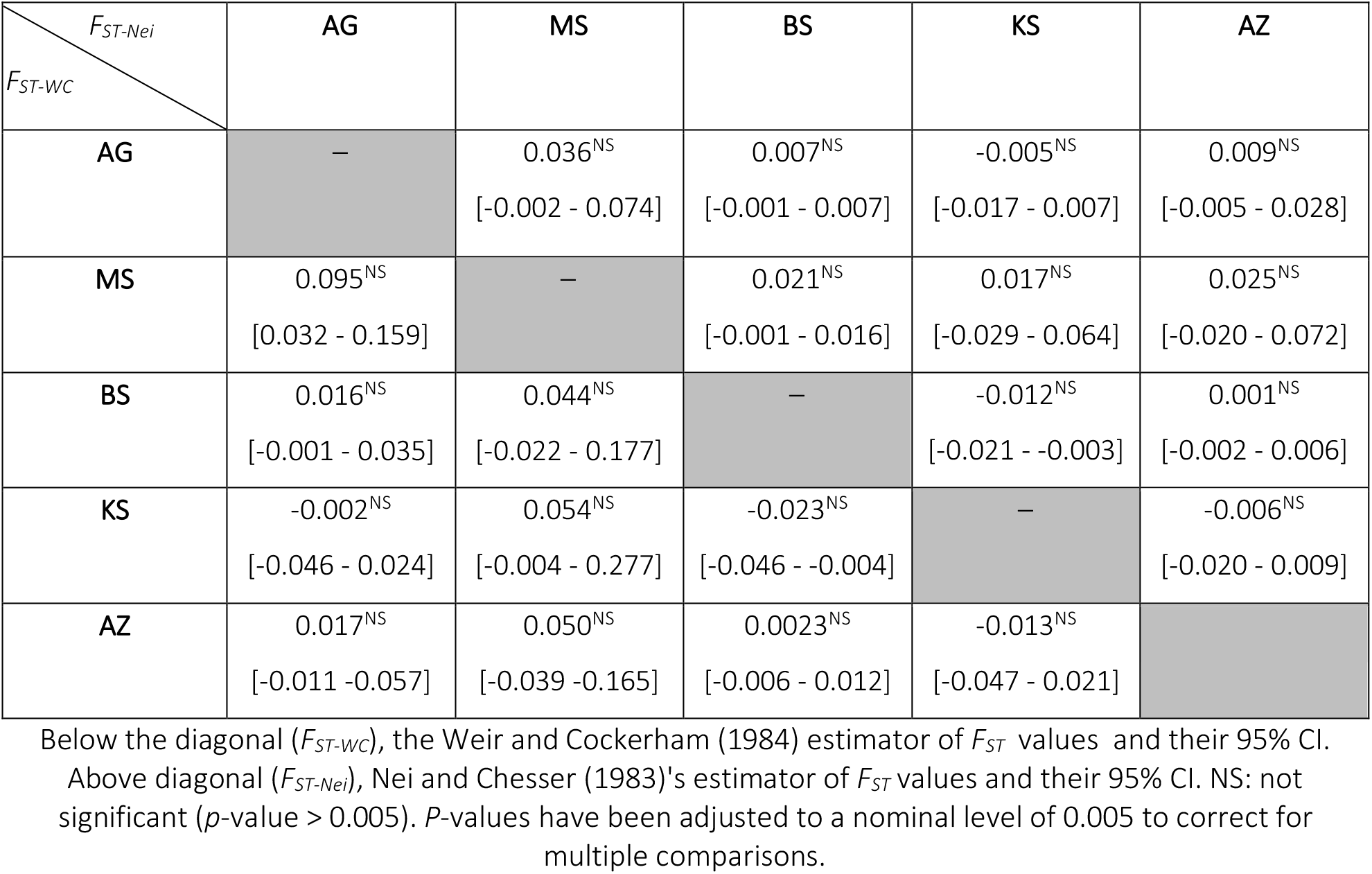
Pairwise *F*_*ST*_ between sampling sites for microsatellites.

The simulation-based assessment of the statistical power to detect significant differentiation as performed in *POWSIM* (Fig. S4) indicated that our microsatellites and mitochondrial datasets have the power to detect significant differentiation for *F*_*ST-Nei*_ > 0.008 and *F*_*ST-Nei*_ >0.1 respectively (Fig. S4). Given the observed Nei’s *F*_*ST*_ values, only the microsatellites data set had thus enough power to detect significant differentiation. Therefore, the lack of genetic differentiation observed at these loci among the five sampled locations does not simply result from a lack of statistical power at the microsatellite loci.

### Simulations of “grey zone” of population differentiation

In agreement with Bailleul *et al.* (2018), *F*_*ST*_ values estimated either from the entire simulated populations or from subsample of 50 individuals were very similar irrespective of the effective size (*Ne*) or the number of migrants exchanged (*Ne*_*cur.*_*m*) (Fig. 3). Simulations show that with a constant contemporary effective size (*Ne*_*cur*_) of 1000 reproducing individuals, the power to detect significant genetic differentiation decreases and the number of generations increases with the number of effective migrants *Ne*_*cur.*_*m*. With less than one migrant per generation (*Ne*_*cur.*_*m* ≤ 1), it takes at most 7 generations to obtain a power of 100% to detect significant *F*_*ST*_ and to reach *F*_*ST*_ values ≥ 0.1 after 700 generations. With 10 migrants per generation (*Ne*_*cur.*_*m* = 10), a high power (>80%) to detect significant *F*_*ST*_ is reached in the 20 first generations, then between 20 and 700 generations, the detection capacity varies between 80% and 100% and the *F*_*ST*_ values vary around 0.017. With high connectivity (*Ne*_*cur.*_*m* = 100) between the two diverging populations, the detection ability stays below 40% during the 700 generations and the simulated *F*_*ST*_ values are lower than 0.002 (Fig. 3). Variation in the initial *Ne* of the founding ancestral population (*Ne*_*ini*_), which mirrored the founding event of the harbor porpoises population in the Black Sea 700 generation ago (Fontaine *et al.*, 2012), had no effect on the detection capacity and on the *F*_*ST*_ values (Fig. 3).

**Fig. 3.**
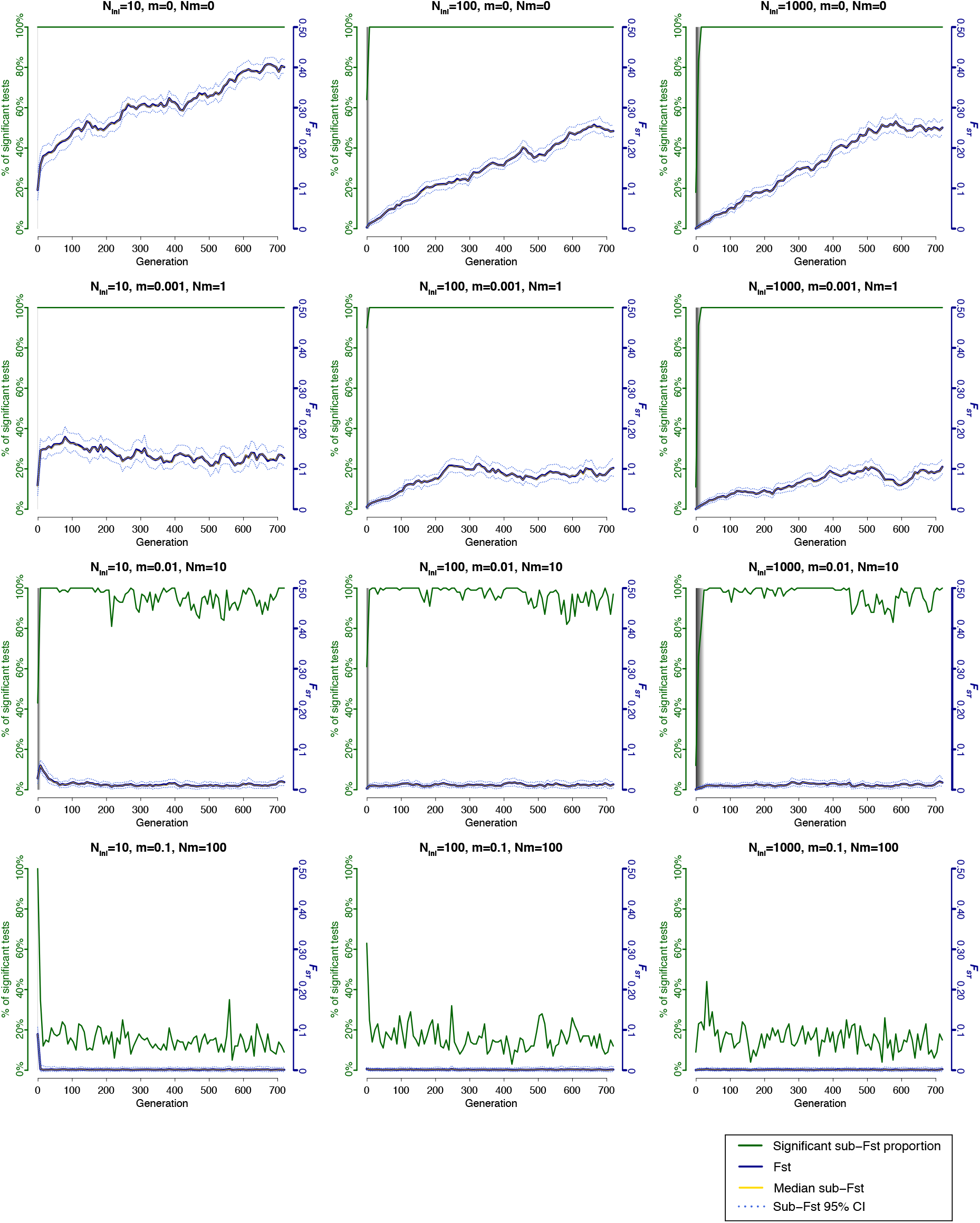
Impact of the “*grey zone*” population differentiation, varying level of connectivity, and number of founders on the genetic differentiation between two hypothetical diverging populations illustrated using simulations. Simulations correspond to two populations, each one with an effective size of 1000, splitting from a small ancestral population with variable initial sizes (*N*_*ini*_), and variable migration rates (*m*) and number of migrants (*N.m*) after the split. For each plot, the x-axis shows the number of generations since the split from the ancestral population, the right y-axis displays the evolution of *F*_*ST*_ values. The median *F*_*ST*_ values and their 95%CI are displayed in blue plain and dashed lines respectively. The median sub-FST values are displayed as yellow lines. The left y-axis shows the proportion of *F*_*ST*_ values significantly different form zero (green line). The vertical grey shades represent the “*grey zone*” of population differentiation defined as the number of generations since the split during which *F*_*ST*_ values are unlikely to be statistically different from 0 in more than 95% of the cases.

## Discussion

### Panmixia in harbor porpoises from the Black Sea and adjacent waters

The widespread genetic homogeneity observed in the Black Sea harbor porpoises is consistent with previous investigations that reported similar results between individuals from the Aegean and Black Seas (Fontaine *et al.*, 2012). Here we report that this homogeneity further extends to the new zones surveyed in this study, including the Crimea, Kerch straight, and the Azov Sea. A genetic panmixia suggests that random mating occurs across the subspecies distribution or that population subdivision is too weak or too recent to have left a detectable signature on the genetic markers analyzed in this study. Such homogeneity is supported by the absence of clustering of the microsatellite genotypes in the *STRUCTURE* (Fig. 2c) and PCA analyses (Fig. 2d), and the absence of significant differences in genetic diversity (Table 1) and allelic frequencies (Table 2). The *POWSIM* power analysis showed that this homogeneity does not result from a lack of power of the microsatellite data to reject panmixia, since simulated datasets with the same number of markers and comparable genetic diversity would be able to detect significant *F*_*ST*_ values as low as 0.008. Mitochondrial data also supported such homogeneity, with no differentiation between the porpoises from the Black and Azov Seas. However, we cannot rule out that the small mitochondrial sample size may reduce the power to reject panmixia.

### High relatedness among porpoises from Marmara Sea biases the population genetic structure

Among the five areas analyzed, three porpoises from the Marmara Sea departed from the others. The DAPC (Fig. S3) and, to a lesser extent, the slightly elevated *F*_*ST*_ values in all the comparisons involving the porpoises from Marmara Sea (Table 2) suggest that the microsatellite allelic frequencies of these porpoises were slightly different from those of the neighboring areas. Yet, only the comparison involving the porpoises from the Marmara and Aegean Seas displayed a *F*_*ST*_ value with a 95% CI excluding the zero. A similar observation was made by Tonay *et al.* (2017) who reported low but statistically significant *F*_*ST*_ values when comparing the mitochondrial haplotype frequencies between the porpoises from the Marmara Sea and those from the Aegean or Black Seas. However, neither Tonay *et al.* (2017)’s results nor ours detected significant *F*_*ST*_ values between the porpoises from the Aegean Sea and Black Sea. From an oceanographic point of view, the Black, Marmara, and Aegean Seas are highly interconnected by strong surface currents flowing from the first into the latter (Aydogdu et al. 2018). Therefore, finding Marmara porpoises genetically differentiated from those of the neighboring seas would be a puzzling result given the intermediate position of the Marmara Sea as a bridge connecting the Aegean Sea and Black Sea. Other factors may explain this result. First, the small sample size in the Marmara Sea (n=3 in our study and n=5 in Tonay et al., 2017) precludes reliable estimation of allele frequency, given the high variance of the stochastic lineage sorting through generation, and especially with such small number of genetic markers (Fogelqvist *et al.*, 2010). Thus, any significant differences in allele frequencies may poorly represent the actual population allele frequencies. Second, a sampling bias that includes closely related individuals can also produce spurious signal of population structure (Anderson and Dunham, 2008; Rodriguez Ramilio and Wang, 2012). For instance, Anderson & Dunham (2008) showed that including individuals from a same family can produce strong spurious signal of population structure within a panmictic population. Average relatedness estimate based on the multilocus microsatellite genotypes showed a very high value in Marmara porpoises (0.55, Table S2 and Fig. S1) consistent with a full siblings or parent-offspring relationship. This elevated relatedness was significantly higher than the observed values in any other localities, and significantly greater than what would be expected by chance (Fig. S1). An elevated relatedness among Marmara porpoises rather than an actual population subdivision likely explains their sight distinctiveness observed at the *F*_*ST*_ values (Table 2) and along the second axis of the DAPC (Fig. S3). The observations of Tonay *et al.* (2017), revealing a subtle but statistically significant mitochondrial differences between porpoises from the Turkish Straights system and the other areas may reflect a similar effect. However, no definitive conclusions can be drawn from the analysis of a single haploid locus. A more robust population and genomic sampling is required to address this question. Nevertheless, assuming that the mitochondrial observations of Tonay *et al.* (2017) do not reflect a relatedness bias, but real population differentiation, the contrast between mitochondrial heterogeneity and microsatellite homogeneity may be consistent with sex-biased dispersal. Females philopatry and males mediating gene flow was reported multiple times in harbor porpoises from North Atlantic waters (e.g. Wang *et al.*, 1996; Rosel *et al.*, 1999).

### An expected homogeneity given the large dispersal abilities of the porpoises

A large-scale genetic panmixia is expected and frequently reported in highly mobile marine species living in an environment where geographical barrier to dispersal are scarce (Quintela *et al.*, 2014; Bailleul *et al.*, 2018). In the case of the Black Sea porpoises, such homogeneity could be expected given the large oceanographic connectivity among the adjacent seas (ex. Aydoğdu *et al.* 2018), the very large dispersal abilities and habitat occupation of the species reported in other areas. For example, Nielsen *et al.* (2018) showed that the total habitat occupation of 72 porpoises tagged in the Danish waters of the North Sea could reach up to ~600,000 km^2^. Even more striking, 30 porpoises from Western Greenland displayed large scale offshore movements and occupied a total habitat of 4,144,749 km^2^. Daily travelling rates can ranged between 20 to 50 km in a single day (Nielsen *et al.*, 2018). Thus, the dispersal abilities of harbor porpoises are comparable or can exceed the total surface of the Black Sea (436, 000km^2^), Azov Sea (39,000 km^2^), Aegean Sea (214,000 km^2^), and Marmara Sea (11,350 km^2^). Furthermore, the continental climate prevailing in the northern Black Sea and Azov Sea can lead to rapid ice formation forcing porpoises to leave the Azov Sea during winter when it becomes completely frozen (Matishov *et al.*, 2014). Massive porpoise mortalities due to ice entrapment have been reported in the past (Kleinenberg, 1956; Birkun, 2002). Therefore, the absence of barriers to gene flow, the large dispersal abilities of the species, the small geographic scale, the frequent movements reported between the different seas (Kleinenberg, 1956; Vishnyakova *et al.*, 2013), and the unavailability of some habitats for part of the year, all points toward highly connected demes of porpoises in the Black Sea and adjacent waters. The simulations (Fig. 3) showed that moderate levels of connectivity (*Ne.m* = 10 migrants per generation) could lead to weak, but still rapidly detectable differences in allelic frequencies in less than 20 generations. Such a result is conservative, since the simulations assumed an effective number of reproducing individuals (*Ne*) of 1000 in each hypothetical diverging group. *Ne* would be probably smaller in nature, leading to an even faster genetic drift of the allele frequencies, and thus to a faster detection ability.

### The ambiguous *“grey zone”* of population differentiation

A number of factors may also limit our ability to detect genetic structure if it exists and warrant a cautionary interpretation of the genetic homogeneity reported in this study, especially for designing conservation strategies. Genetic homogeneity does not necessarily imply demographic homogeneity. A certain number of generations after a population split is required for genetic drift to change the allele frequencies in diverging populations and reach the migration-drift equilibrium (Epps and Keyghobadi, 2015). The time-lag before which genetic variation become a good proxy of demographic subdivision is dependent on *Ne* and thus on the life history traits of the species (Waples, 1998; Gagnaire *et al.*, 2015; Bailleul *et al.*, 2018). In species exhibiting high fecundity and large population sizes (e.g. fish such as herring, anchovy, salmon, blue shark) genetic drift can be ineffective and genetic differentiation very weak or even absent (Waples, 1998; Gagnaire *et al.*, 2015; Bailleul *et al.*, 2018). In those species, a lack of genetic differentiation can therefore result from a range of situations spanning from nearly complete demographic independence among large-sized populations to the existence of a unique panmictic population (Palumbi, 2003; Gagnaire *et al.*, 2015). In contrast, cetacean species, like porpoises, display much more reduced fecundity and population sizes (Hoelzel, 1998; Read 1999). In those species, simulations show that the *“grey zone”* of population differentiation is narrow (Fig. 3). Rejection of the panmixia can be quickly achieved with good power even for moderate level of gene flow (*Ne.m*=10 or 1% of the total population size). Such power vanishes when connectivity reaches 10% of the total population size (*Ne.m* = 100 migrants per generation). Under such high connectivity, our simulations are consistent with previous works that shows that the level of migration rate no longer allow demographic units to be independent (Palsbøll *et al.*, 2007). Nevertheless, under a moderate level of gene flow (*Ne.m=*10 or 1%), rejection of panmixia is achieve with good power in less than 20 generations. In the case of the harbor porpoise, this corresponds to 200 years (assuming a conservative generation time of 10 years, Read 1999). This means that in most case populations that are isolated enough would be detectable with relatively good power, unless the split occurred within the last 20 generations.

### More robust dataset and sampling design may reveal cryptic structure

Both empirical and simulation-based studies shows that massively increasing the number of loci genotyped across the genome can improve the power to detect subtle population structure when it exists (Willing *et al.*, 2012; Funk *et al.*, 2012). Thousands of single nucleotide polymorphisms (SNPs) can now be obtained at a reasonable cost using reduced representation of genomic variation (e.g., RADseq, GBS) (Davey *et al.*, 2011) or low coverage whole-genome sequencing (Cammen *et al.*, 2016). Genomic data set may outperform microsatellite and mitochondrial markers to reveal subtle population subdivisions (Funk *et al.*, 2012). Not only increasing the number of loci would offer a more representative picture of the genomic variation, SNPs genotyping could also deliver more accurate genotyping than microsatellite loci (Morin *et al.*, 2004; Willing *et al.*, 2012). In situations where population structure is weak, as it may be the case for the Black Sea porpoises, analyzing thousands of SNPs may reveal subtle patterns of populations structure. This has been illustrated recently in the case of harbor porpoises from the Baltic Sea. The use of ~3000 SNP loci revealed subtle population subdivisions between porpoises from the Baltic and North Seas that could not be accurately resolved using microsatellite and mitochondrial loci (Lah *et al.*, 2016).

Beyond the sample size, number of loci, and a possible ambiguous population “*grey zone”*, numerous species, including porpoises, exhibit specific breeding behaviors, aggregating only during the reproductive season. Grouping behaviors were recently identified in the Baltic Sea during the reproductive season that could possibly reflect breeding groups (Carlén *et al.*, 2018). Field surveys in the Kerch Strait and in the north-eastern Black Sea also suggested that such aggregating behavior may exists (Vishnyakova *et al.*, 2013). Although most of the sampling used in this study are from animals stranded during the breeding season (Gol’din, *pers. comm.* and Vishnyakova and Gol’din 2015), we cannot rule out that aggregating samples over seasons and years may limit the power to discover subtle population structure if it exists (Wiemann *et al.*, 2010; Bailleul *et al.*, 2018; Graves and McDowell, 2015). Comparing groups defined *a priori* based on subjective delimitations (such as geography), neglecting seasonal and yearly effects, is fraught with power limitations. Exploratory individual-based approaches, such as principal component analyses (Jombart *et al.*, 2009) and Bayesian clustering analyses (Pritchard *et al.*, 2000), as used in this study, may partly compensate for this limitation (Fontaine *et al.*, 2007; Guillot *et al.*, 2009). However, the statistical power of these approaches depends on the sample size, the number of genetic markers, and the magnitude of population structuration (Morin *et al.*, 2004; Willing *et al.*, 2012). A sampling design providing the flexibility to contrast sex and seasonal effects may reveal subtle pattern of population structure that would not be amenable otherwise. For example, by stratifying the samples according to sample type (stranding vs. by-caught), sex, and season (breeding vs. non-breeding season), Wiemann *et al.* (2010) were able to provide the first evidence of population structuration in the harbor porpoises from the Baltic Sea using microsatellite markers that were later confirmed with genome-scale data (Lah *et al.*, 2016).

### A genetic homogeneity in face of morphological heterogeneity

Significant morphological differences were previously reported between the porpoises from the Black Sea and Azov Sea (Gol’din, 2004; Gol’din and Vishnyakova, 2015; 2016). These authors hypothesized that such phenotypic differences could reflect demographically, ecologically and genetically differentiated groups. However, our results currently do not support this hypothesis. Such a discrepancy between genetics and morphology have been widely reported (Rheindt *et al.*, 2011). A first plausible explanation could be that the observed morphological variations between Azov and Black Sea porpoises are related to epigenetic variation. Epigenetic mechanisms, which regulate the expression of genes without modifying the DNA, can induce phenotypic plasticity when exposed to different ecological conditions (Duncan *et al.*, 2014). If porpoises are adapted to distinct cryptic local environmental conditions, morphological differences could result from such phenotypic plasticity, without being underpinned by genetic variation. A second plausible explanation is that the handful selectively neutral loci used in this study may not reveal genetic differentiation occurring in other places of the genome that are involved in this ecological adaptation (Gagnaire *et al.*, 2015). Markers that depart from neutral expectations can form localized island of differentiation along the genome and are good evidence that divergent adaptive processes are ongoing (Turner and Hahn, 2010). Examples of such genomic islands of differentiation have been reported in sticklebacks (Ravinet *et al.*, 2018), cichlid fishes (Malinsky *et al.*, 2015), and Anopheles mosquitoes (Turner and Hahn, 2010). They are characteristic of incipient ecological differentiation. In porpoises, phenotypic variation may be related to natural selection happening in portions of the genome that are not targeted by the neutral markers used in this study. Whole genome sequencing data are required and may reveal cryptic differentiation. Revealing these patterns could be of paramount importance since the signal held by outlier loci could be useful to delineate locally adapted stocks and conservation units (Funk *et al.*, 2012; Gagnaire *et al.*, 2015).

## Conclusions

Despite previous evidence of phenotypic heterogeneity between the porpoises from the Azov and Black Seas, the present study did not allow to reject the panmixia hypothesis. This result does not necessarily imply a population homogeneity since a number of factors may limit our ability to detect subtle pattern of population subdivision. The limited sample size at the population and genomic level, and the neutral loci used here may have missed cryptic differentiation. Furthermore, we cannot rule out that population structure may be too weak or too recent to be capture with neutral genetic loci. Therefore, this study stresses the need to improve the sampling design in order to maximize the power of the analyses to reject panmixia in all areas where the species occurs and to provide meaningful contrasts that can impact the power to detect subtle population structure if it exists (e.g. sex, season and age class). Next Generation Sequencing approaches to increase massively the number genetic markers along the genome could improve the statistical power to detect subtle population structure and provide means to address hypotheses of ecologically differentiated groups in the Black Sea porpoises that could only be observable at specific loci in the genome that are impacted by natural selection.

From a conservation perspective, the recent report of the status of harbor porpoise in the North Atlantic workshop by the NAMMCO/IMR (2019) recently reiterated the crucial need to define management units based on the total amount of evidence available to assess the threats and manage the species in the areas under focus. Improving population genetics information will be needed for better understanding the population structure and connectivity among porpoises in the Black Sea and adjacent waters. In addition, sustained efforts should continue to gather additional information on the variation in ecology, morphology, demography, life history traits and other biological meaningful metrics, as well as on the stranding and by-catch rates to make informed management decisions.

## Acknowledgments

We thank Diane Bailleul for sharing the scripts to perform the simulations and Jan Veldsing for his assistance in the laboratory. We are also grateful to Jeanine Olsen for her feedbacks that greatly improved the manuscript. This study benefitted from funding of the University of Groningen (The Netherlands) through a PhD fellowship allocated to YBC. We would like to thank the Center for Information Technology of the University of Groningen for their support and for providing access to the *Peregrine* high-performance computing cluster.

## Author Contributions

MCF and YBC designed the study; KV and PG collected the samples in the field; JT performed the laboratory work and collected the data; YBC and MCF analyzed the data; YBC and MCF interpreted the results and wrote the manuscript with input and final approval from all the co-authors.

## Data archiving

Data and scripts available from the Dryad Digital Repository: (link to be announced)

