## Supplementary Materials for "Genetic homogeneity in face of morphological heterogeneity in the harbor porpoises from the Black Sea and adjacent waters"

Content

**Table S1 |** Nuclear microsatellite genetic diversity for each population.

**Table S2 |** Within and between population relatedness.

**Fig. S1 |** Analysis of relatedness within groups.

**Fig. S2 |** Population structure estimated using *STRUCTURE*.

**Fig. S3 |** Scatter plot showing the first two discriminant functions of the DAPC.

**Fig. S4 |** *POWSIM* statistical assessment of the power to detect significant differentiation between two populations from microsatellites and mitochondrial datasets.

**Table S1.** Nuclear microsatellite genetic diversity for each population

| | | EV94 | GT011 | igf-1 | PPH104 | PPH110 | PPH130 | 415-416 | GATA053 | GT015 | PPH137 | All |
|---|---|---|---|---|---|---|---|---|---|---|---|---|
| AG | *n* | 11 | 10 | 11 | 9 | 11 | 11 | 0 | 0 | 9 | 9 | 9.2 |
| | *Ar*[1] | 1.48 | 1.33 | 1.84 | 1.77 | 1.59 | 1.62 | NA | NA | 1.47 | 1.45 | 1.51±0.08 |
| | *pAr*[1] | 0.01 | 0.04 | 0.61 | 0.58 | 0.37 | 0.1 | NA | NA | 0.19 | 0.04 | 0.22±0.08 |
| | $H_o$ | 0.55 | 0.40 | 0.73 | 1.00 | 0.73 | 0.91 | NA | NA | 0.44 | 0.44 | 0.58 |
| | *He* | 0.46 | 0.32 | 0.81 | 0.73 | 0.57 | 0.60 | NA | NA | 0.44 | 0.43 | 0.54 |
| | $F_{IS}$ | -0.18$^{NS}$ | -0.25$^{NS}$ | 0.10$^{NS}$ | -0.37$^{NS}$ | -0.28$^{NS}$ | -0.52$^{NS}$ | NA | NA | 0.00$^{NS}$ | -0.04$^{NS}$ | -0.19$^{NS}$ |
| MS | *n* | 3 | 3 | 3 | 3 | 1 | 3 | 2 | 3 | 3 | 3 | 2.7 |
| | *Ar*[1] | 1.60 | NA | 1.60 | 1.73 | 1.33 | 1.60 | NA | NA | 1.33 | 1.53 | 1.49±0.1 |
| | *pAr* | 0.02 | NA | 0.77 | 0.44 | 0.96 | 0.28 | NA | NA | 0.07 | 0.13 | 0.30±0.11 |
| | $H_o$ | 1.00 | NA | 1.00 | 1.00 | 1.00 | 0.33 | 0.50 | NA | 0.33 | 0.67 | 0.58 |
| | *He* | 0.50 | NA | 0.50 | 0.61 | 0.50 | 0.50 | 0.38 | NA | 0.28 | 0.44 | 0.37 |
| | $F_{IS}$ | -1.00$^{NS}$ | NA | -1.00$^{NS}$ | 0.64$^{NS}$ | -1.00$^{NS}$ | 0.33$^{NS}$ | -0.33$^{NS}$ | NA | -0.20$^{NS}$ | -0.50$^{NS}$ | -0.54$^{NS}$ |
| BS | *n* | 86 | 86 | 86 | 86 | 84 | 87 | 79 | 86 | 83 | 86 | 84.9 |
| | *Ar*[1] | 1.49 | 1.43 | 1.75 | 1.65 | 1.51 | 1.65 | 1.44 | 1.02 | 1.35 | 1.66 | 1.49±0.07 |
| | *pAr*[1] | 0.08 | 0.11 | 0.38 | 0.36 | 0.18 | 0.21 | NA | 0.02 | 0.29 | 0.26 | 0.21±0.04 |
| | $H_o$ | 0.48 | 0.48 | 0.78 | 0.66 | 0.48 | 0.59 | 0.37 | 0.02 | 0.39 | 0.73 | 0.50 |
| | *He* | 0.48 | 0.43 | 0.74 | 0.65 | 0.50 | 0.64 | 0.44 | 0.02 | 0.35 | 0.66 | 0.49 |
| | $F_{IS}$ | 0.01$^{NS}$ | -0.11$^{NS}$ | -0.05$^{NS}$ | -0.03$^{NS}$ | 0.06$^{NS}$ | 0.09$^{NS}$ | 0.16$^{NS}$ | -0.01$^{NS}$ | -0.10$^{NS}$ | -0.12$^{NS}$ | -0.01$^{NS}$ |
| KS | *n* | 6 | 7 | 7 | 7 | 7 | 7 | 3 | 6 | 5 | 7 | 6.2 |
| | *Ar*[1] | 1.53 | 1.53 | 1.79 | 1.79 | 1.44 | 1.73 | 1.53 | NA | 1.36 | 1.63 | 1.53±0.07 |
| | *pAr*[1] | 0.01 | 0.16 | 0.38 | 0.72 | 0.09 | 0.30 | NA | NA | 0.09 | 0.14 | 0.21±0.08 |
| | $H_o$ | 0.50 | 0.57 | 0.86 | 1.00 | 0.29 | 0.71 | 0.67 | NA | 0.40 | 0.86 | 0.59 |
| | *He* | 0.49 | 0.49 | 0.73 | 0.73 | 0.41 | 0.67 | 0.44 | NA | 0.32 | 0.58 | 0.49 |
| | $F_{IS}$ | -0.03$^{NS}$ | -0.17$^{NS}$ | -0.17$^{NS}$ | -0.36$^{NS}$ | 0.30$^{NS}$ | -0.06$^{NS}$ | -0.50$^{NS}$ | NA | -0.25$^{NS}$ | -0.47$^{NS}$ | -0.19$^{NS}$ |
| AZ | *n* | 29 | 32 | 31 | 30 | 32 | 29 | 14 | 28 | 17 | 31 | 27.3 |
| | *Ar*[1] | 1.55 | 1.31 | 1.76 | 1.72 | 1.54 | 1.65 | 1.47 | NA | 1.17 | 1.69 | 1.49±0.08 |
| | *pAr*[1] | 0.16 | 0.06 | 0.44 | 0.47 | 0.18 | 0.16 | NA | NA | 0.13 | 0.36 | 0.22±0.08 |
| | $H_o$ | 0.52 | 0.31 | 0.68 | 0.70 | 0.66 | 0.38 | 0.36 | NA | 0.18 | 0.71 | 0.45 |
| | *He* | 0.54 | 0.31 | 0.75 | 0.71 | 0.54 | 0.64 | 0.46 | NA | 0.16 | 0.68 | 0.48 |
| | $F_{IS}$ | 0.05$^{NS}$ | -0.01$^{NS}$ | 0.09$^{NS}$ | 0.02$^{NS}$ | -0.22$^{NS}$ | 0.41*** | 0.22$^{NS}$ | NA | -0.07$^{NS}$ | -0.04$^{NS}$ | 0.05$^{NS}$ |

n : sample size ; *Ar*±SD : Allelic richness ; *pAr*±SD : Private allelic richness : $H_o$ : observed heterozygosity ;$H_e$ : expected heterozygosity ; $F_{IS}$ : fixation index of Weir and Cockerham (Weir and Cockerham, 1984); *** : p<0,001; NS : Not significant; [1]Standardized for a sample size of 2 individuals (Szpiech *et al.*, 2008). AG, Aegean Sea; MS, Marmara Sea; BS, Black Sea; KS, Kerch Strait; AZ, Azov Sea.

Table S2: Within and between population relatedness

| | AG | MS | BS | KS | AZ |
|---|---|---|---|---|---|
| **AG** | 0.08±0.31 | - | - | - | - |
| **MS** | -0.04±0.31 | **0.55±0.15*** ** | - | - | - |
| **BS** | 0.01±0.31 | 0.03±0.25 | 0.01±0.28 | - | - |
| **KS** | 0.06±0.34 | 0.05±0.26 | -0.04±0.28 | 0.05±0.30 | - |
| **AZ** | 0.01±0.31 | -0.06±0.27 | -0.03±0.29 | -0.01±0.30 | -0.05±0.30 |

Average ± standard deviation of the coefficient of relatedness Wang (2002) estimator of relatedness among populations (lower matrix) and within populations (along the diagonal) estimated with *Related* v.1.0 (Pew *et al.*, 2015). Significance level estimated using 1000 permutations (*** : p<0,001, Fig. S1). The meaning of the group acronyms is provided in Table S1.

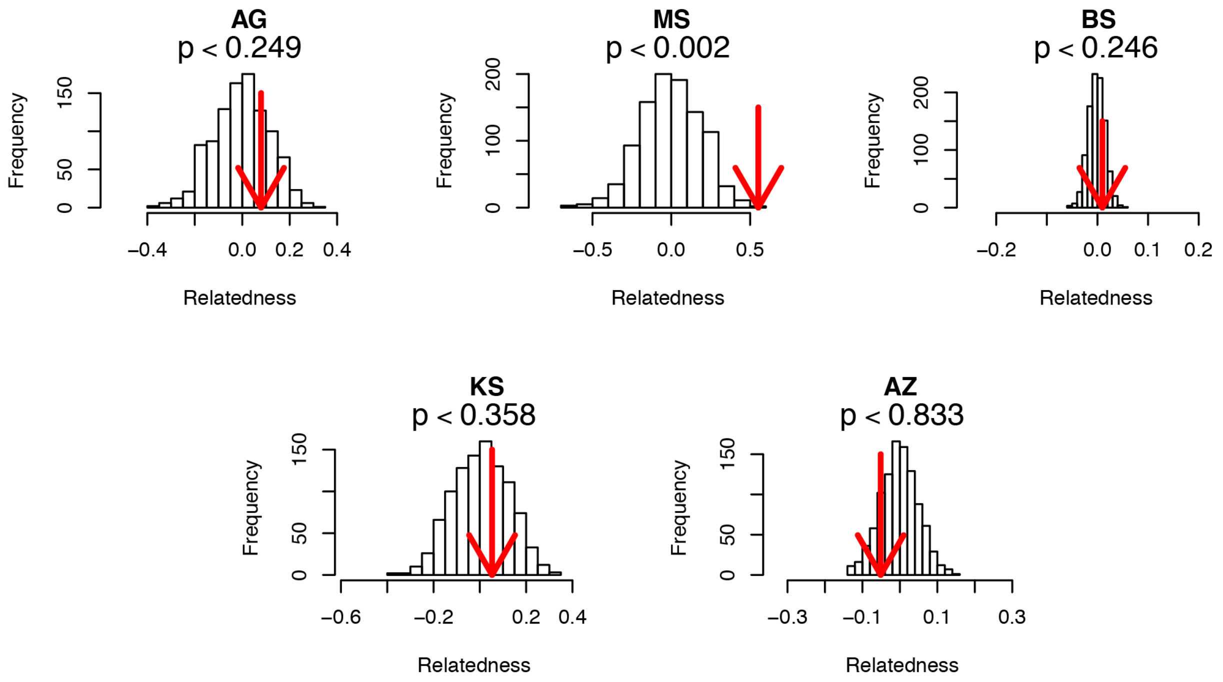


**Fig. S1.** Analysis of relatedness within groups. Each histogram represents the expected Wang (2002) *r* estimator of relatedness within each group. The red arrow shows the observed *r* value for each group in the null distribution expected by chance only. The p-value indicates the proportion of permutations greater than the observed value based on 1000 permutations. The meaning of the group acronyms is provided in Table S1.

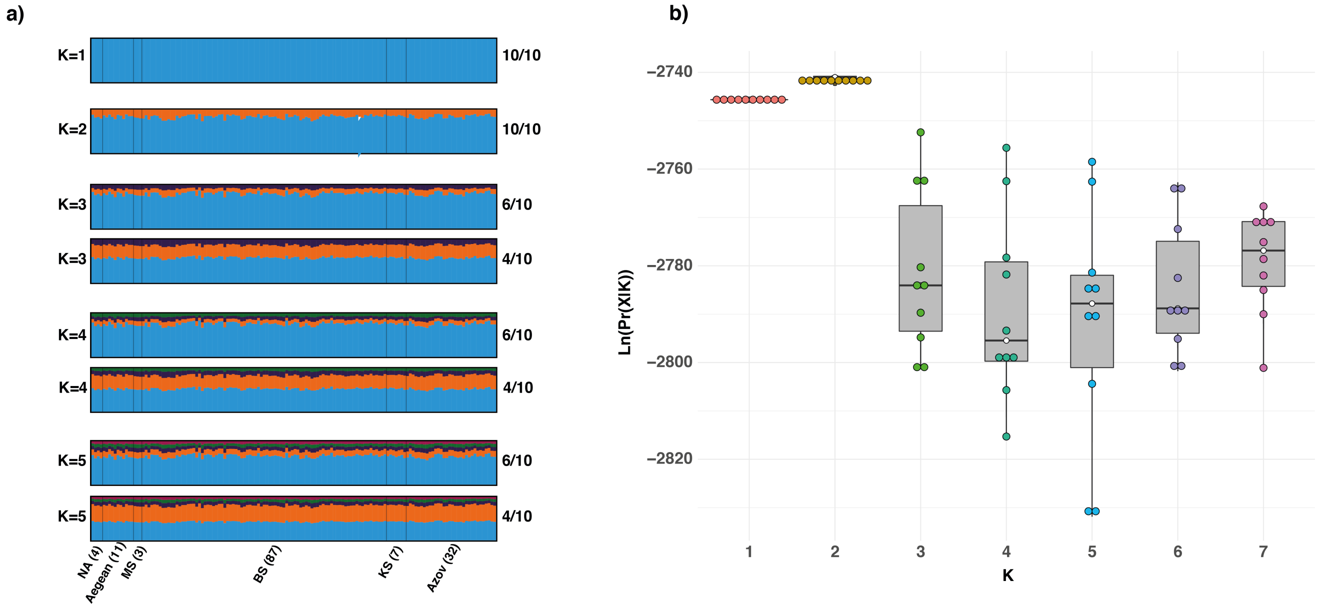


**Fig. S2.** Population structure estimated using *STRUCTURE*. a) Barplots of the individual admixture proportion estimated for K= 1 to K=5. Numbers on the right side of the barplot show the number of times this solution was found out of the 10 runs. b) Distribution of the estimated probability of the data (X) given K groups tested.

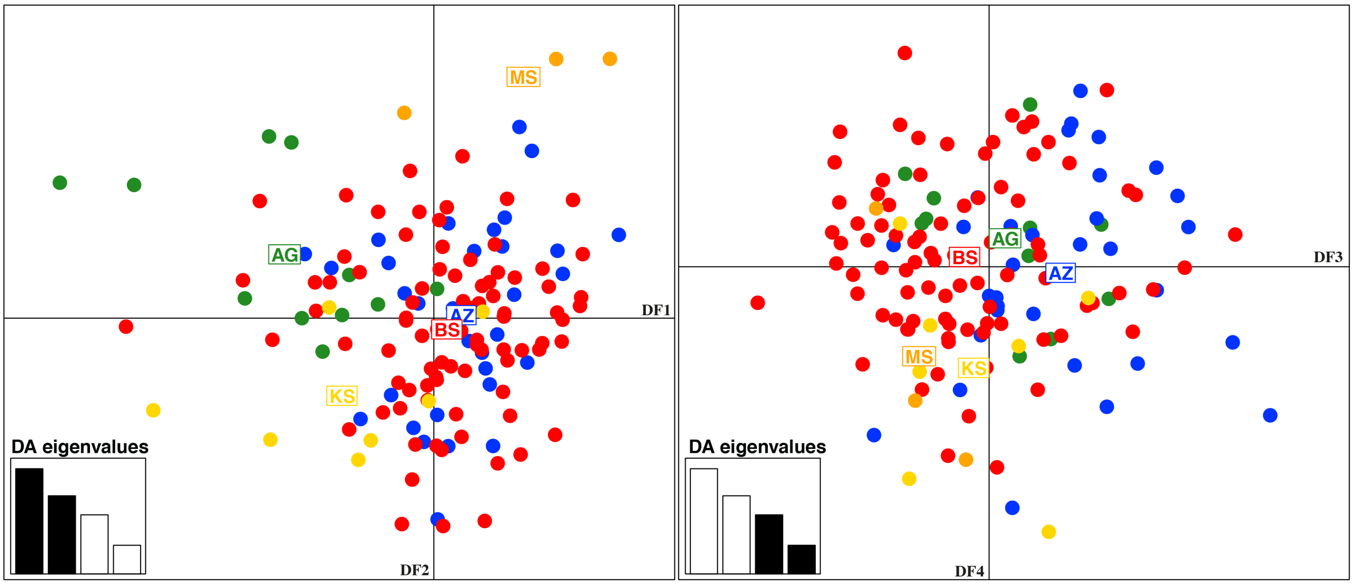


**Fig. S3.** Scatter plot showing the first two discriminant functions of the DAPC. DFs, Discriminant functions; DA, Discriminant analysis. The meaning of the group acronyms is provided in Table S1.

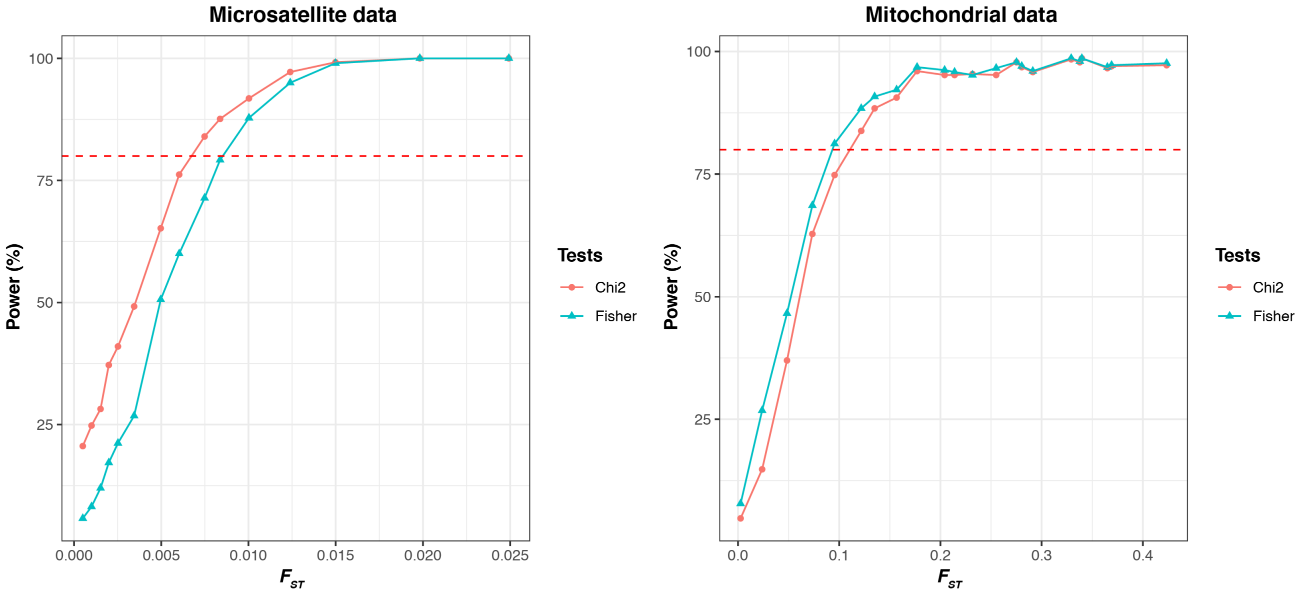


Fig. S4. *POWSIM* statistical assessment of the power to detect significant differentiation between two populations based on the microsatellites (a) and mitochondrial (b) datasets. The dashed line in red represents the power threshold of 80% recommended by Ryman and Palm (2006). *POWSIM*'s results indicate that the microsatellites dataset has the power to detect correctly $F_{ST}$ values from approximately 0.008 and the mitochondrial data set $F_{ST}$ values from approximately 0.1.
